## Supplemental figures for "Order among chaos: high throughput MYCroplanters can distinguish interacting drivers of host infection in a highly stochastic system"

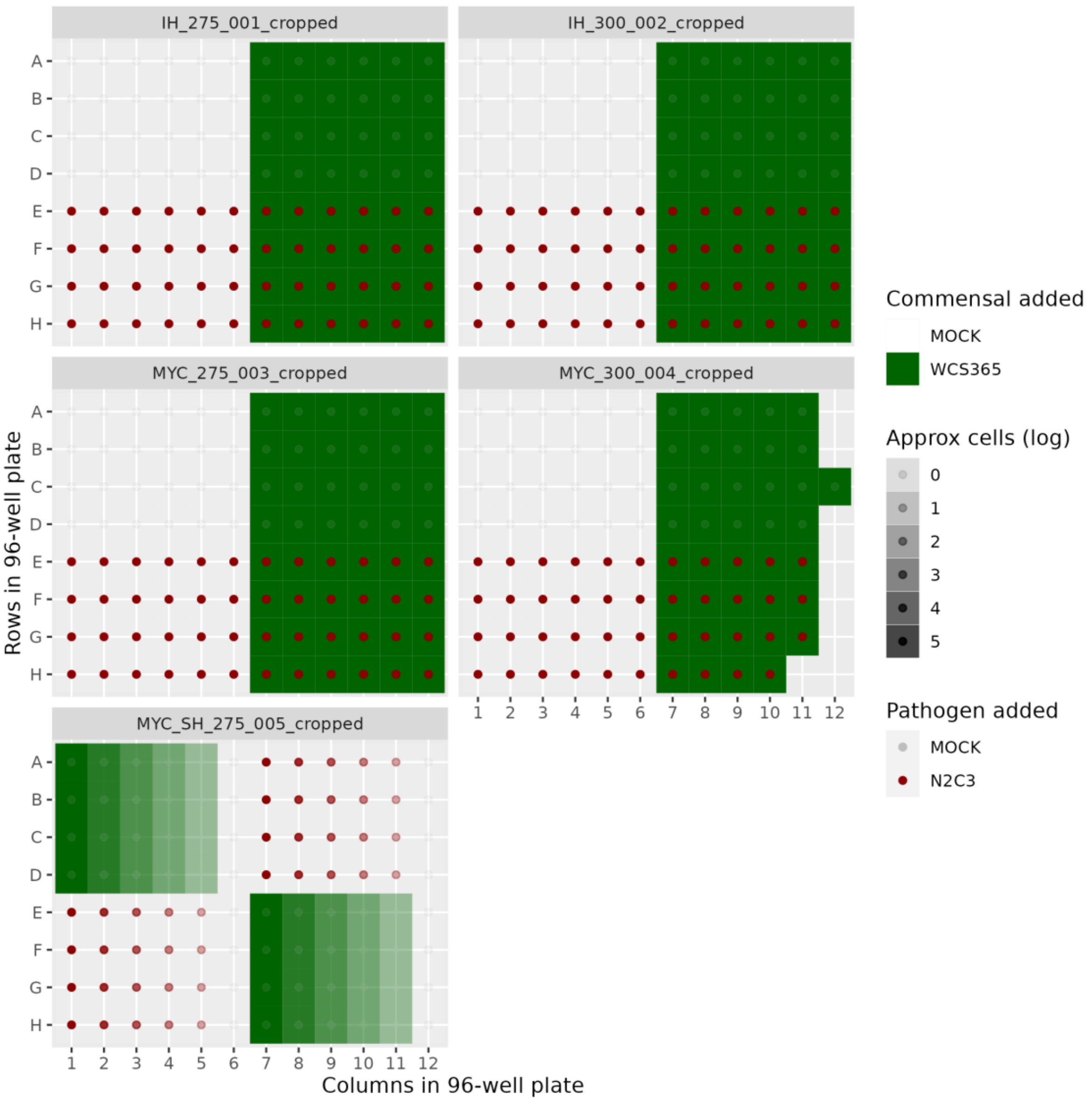


Figure S1. Platemap for N2C3 vs WCS365 pilot experiment to validate MYCroplanter sensitivity in detecting our model system effects. Mock treatments were plain 1/2MS 1/2MES pH5.8 plant growth media.

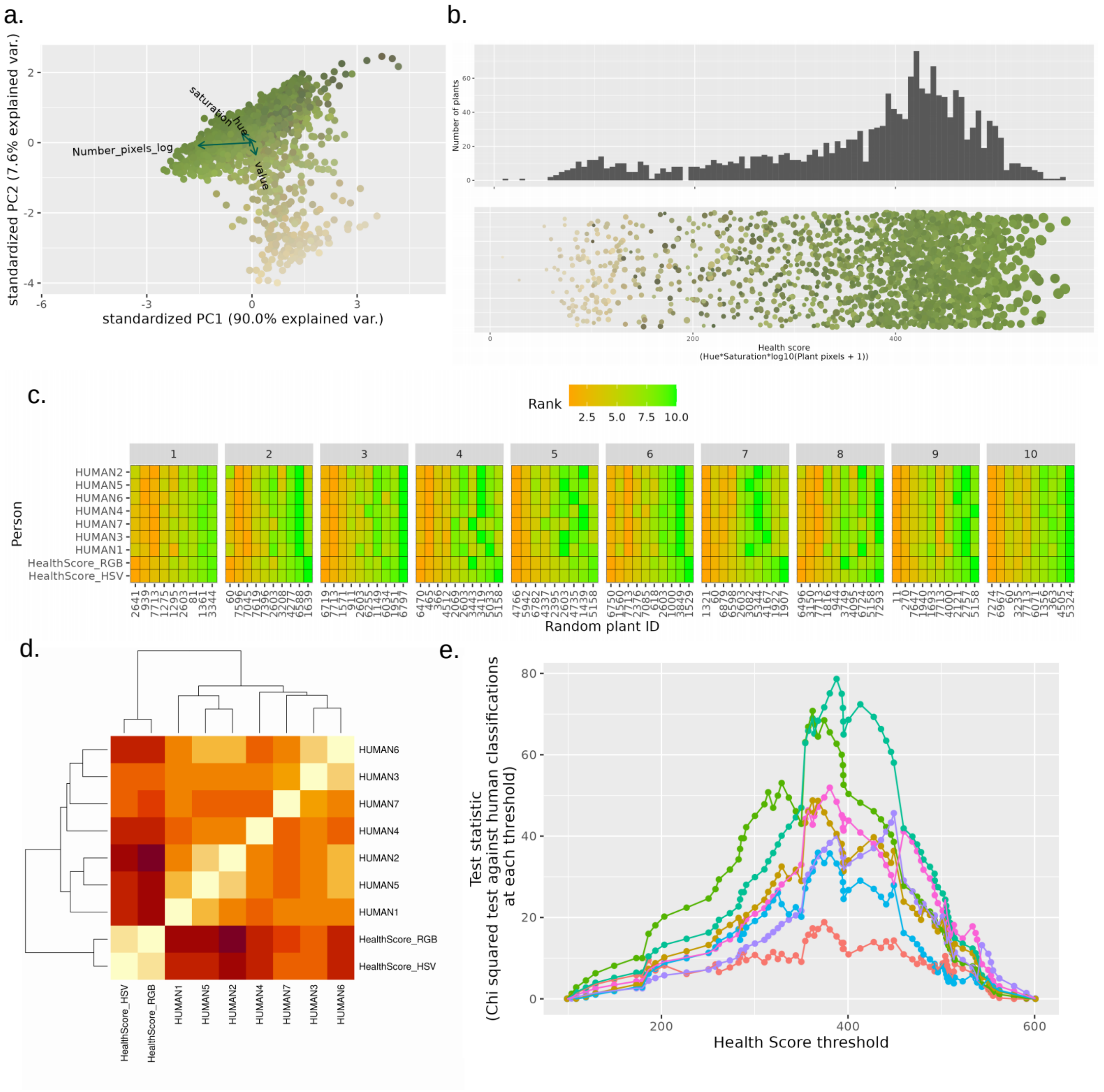


Figure S2. Pixel data from scanned plant images reveals a multi-model distribution of plant health scores that align with human-derived rankings of plant health. (a) Principal components analysis of median pixel values using the HSV (Hue, Saturation, Value) color model. Plants are separated by two primary axis: light green/dark green(healthy/stressed) and green/yellow (alive/dead). (b) HSV pixel values were transformed into a single continuous health score metric (hue*saturation*log10(all_plant_pixels+1)*1000), which reflects the roughly tri-modal distribution of plant health states (healthy / stressed / dead). (c) Algorithm-derived “Health Score” metrics using HSV and RGB values were compared to human-derived ranks. Human rankers ranged from lay-people (no scientific background) to expert (PhD). Humans and scoring algorithms were asked to rank 10 sets of 10 plants (85 total plant images; some images were repeated between sets for validation purposes) in order from “least healthy” to “most healthy”. Each was also asked to divide plants into “not healthy” and “healthy” groups. We found that three plants were consistently ranked differently between humans and our algorithm (7714, 3443, 5158), and that these plants were two-toned (half green half yellow). This caused our algorithm results to perform poorly. When these plants were removed, our Health Score algorithm performed comparably to most human rankers (c). “Healthy” / “Not healthy” classifications by humans were compared to algorithm classifications at different Health Score thresholds. To avoid over-interpretation of health score values, we categorized plants as “healthy” or “not healthy” based at a threshold of 400. Binary response variables are used and reported in the main results and text.

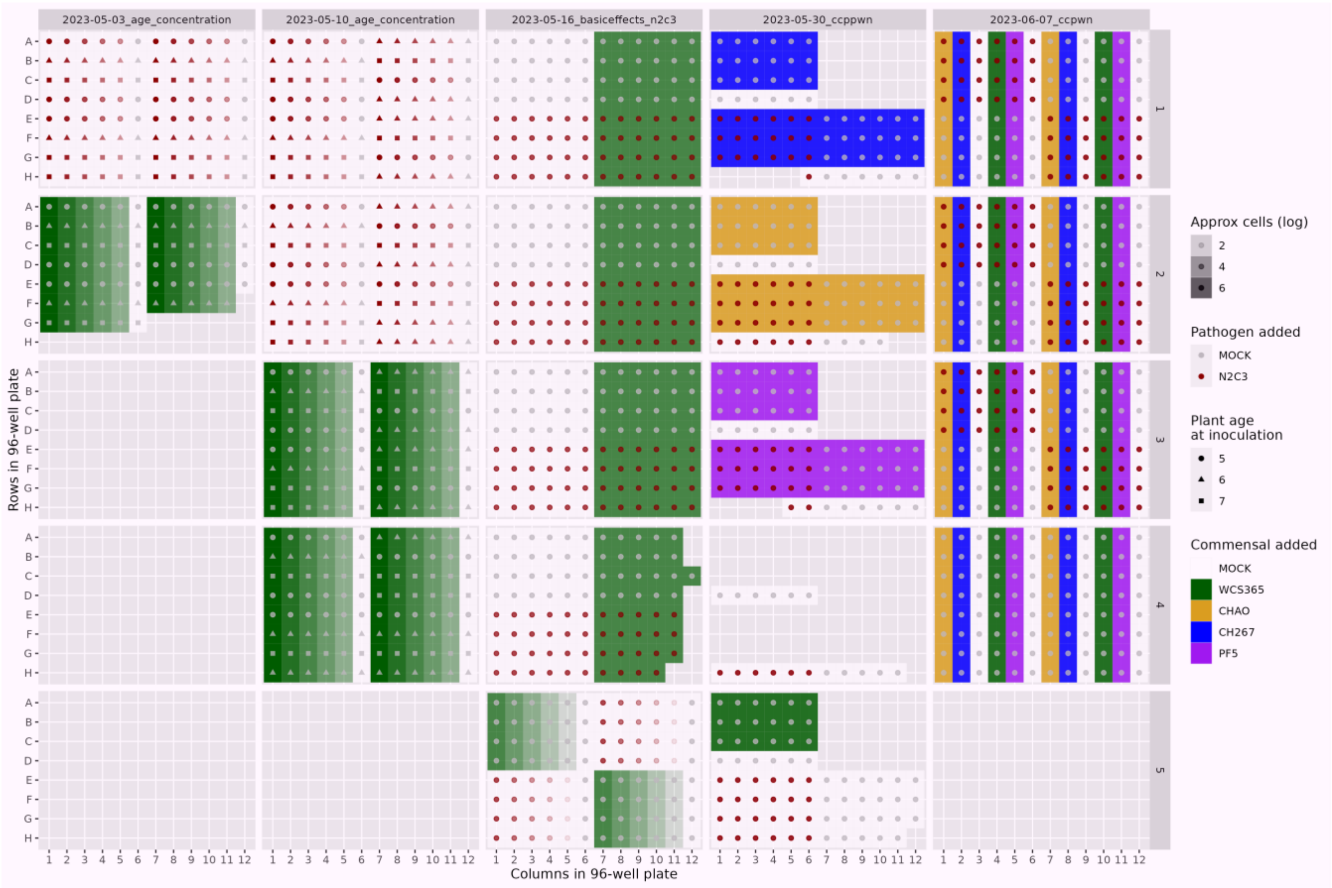


Figure S3. Plate map and experimental design for pilot data used to derive plant health scores. Plants were inoculated at three different ages (5, 6, 7 days) with four different concentrations of pathogen and 4 types of commensal strains across 5 experiments. The bread of experimental variables used was to increase the range of plant health outcomes observed in our pilot data.

a. b.

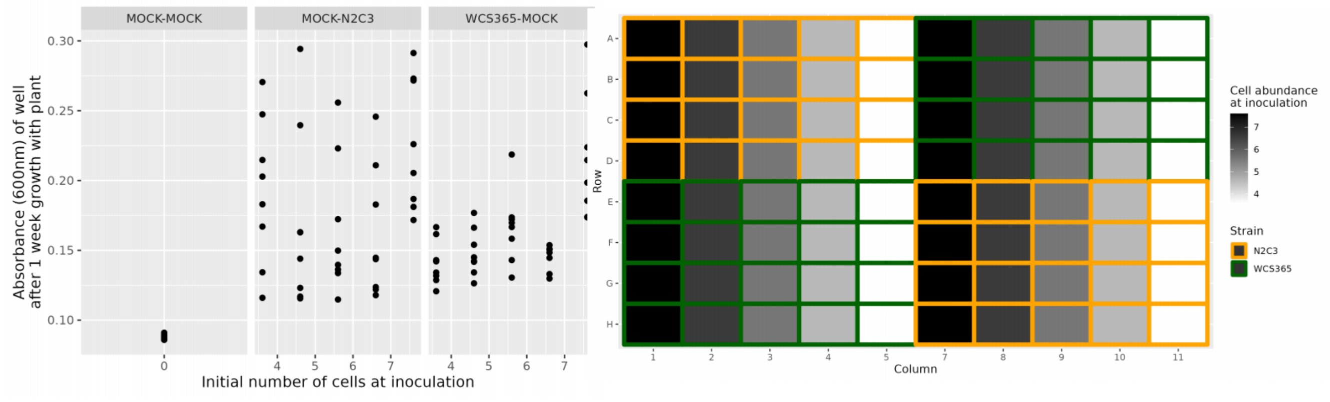


Figure S4. Final absorbance of cells in 96-well plates after 1 week of incubation with plants shows that carrying capacity is similar regardless of initial inoculation concentration. (b) Platemap and experimental design for data shown in (a). Plants were inoculated with different concentrations of pathogen and protective.

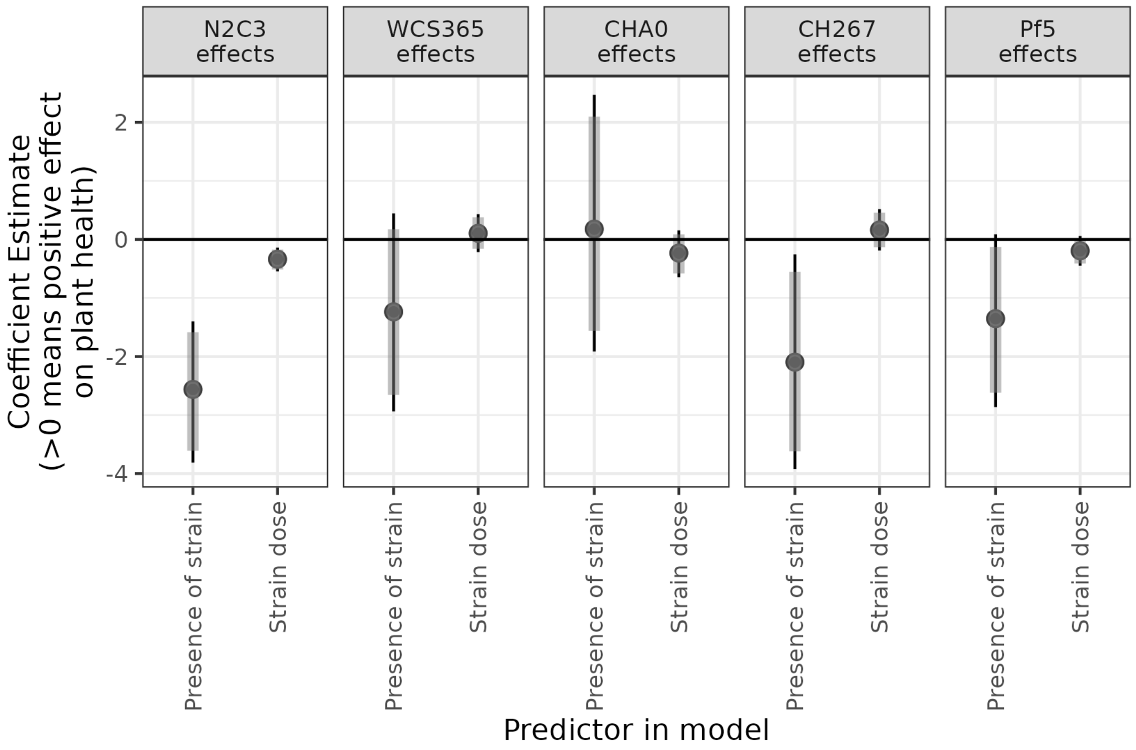


Figure S5. Effect of strain presence and dose for *Pseudomonas* strains in monoculture on probability of plants remaining healthy. Points are median estimates from a Bayesian Bernoulli model, thick bars are 90% credible intervals, and thin bars are 95% credible intervals.

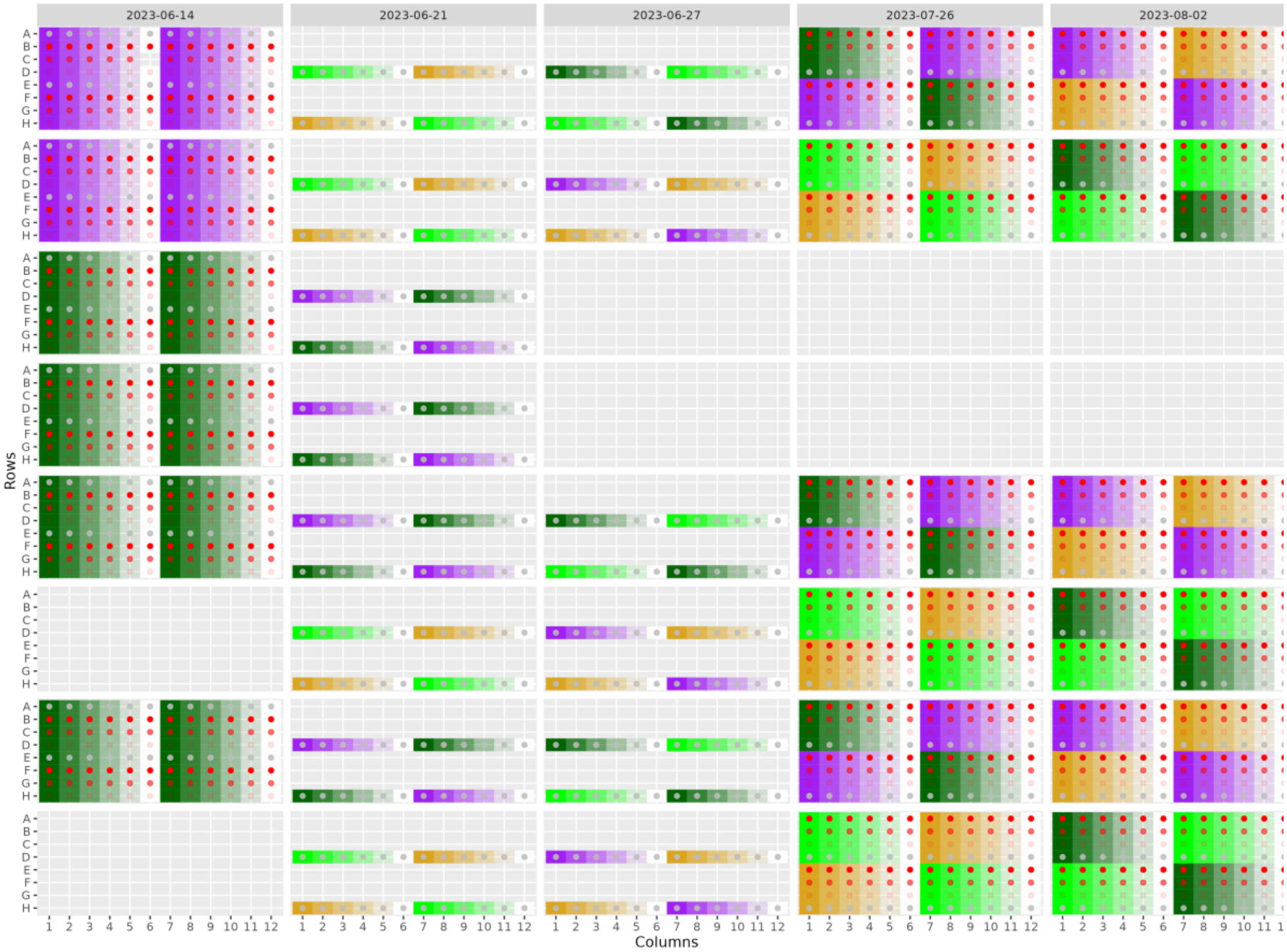


Figure S6. Plate map and experimental design for experiment testing effect of inoculation ratio and plant age on infection outcome. Plants were inoculated at three different ages (5, 6, 7 days) with 4 or 5 concentrations of pathogen and protective across 5 experiments.

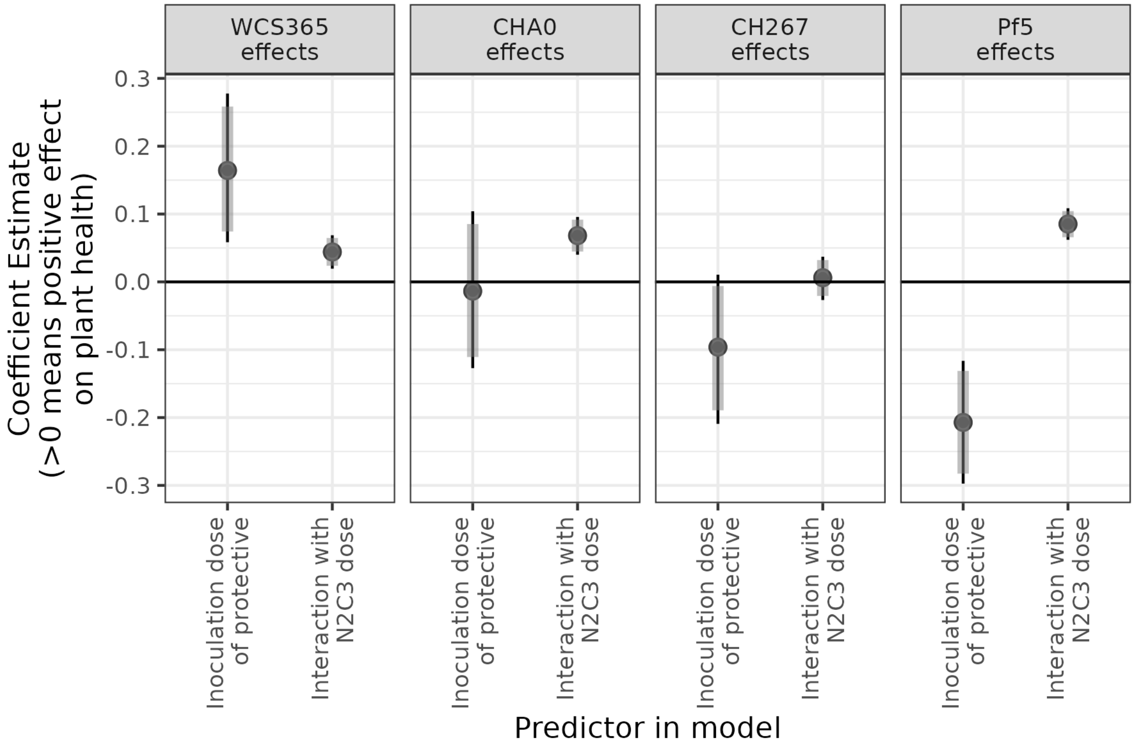


Figure S7. Effect of microbiota and pathogen dose across microbiota genotype on probability of plants remaining healthy. Points are median estimates from a Bayesian Bernoulli model, thick bars are 90% credible intervals, and thin bars are 95% credible intervals.

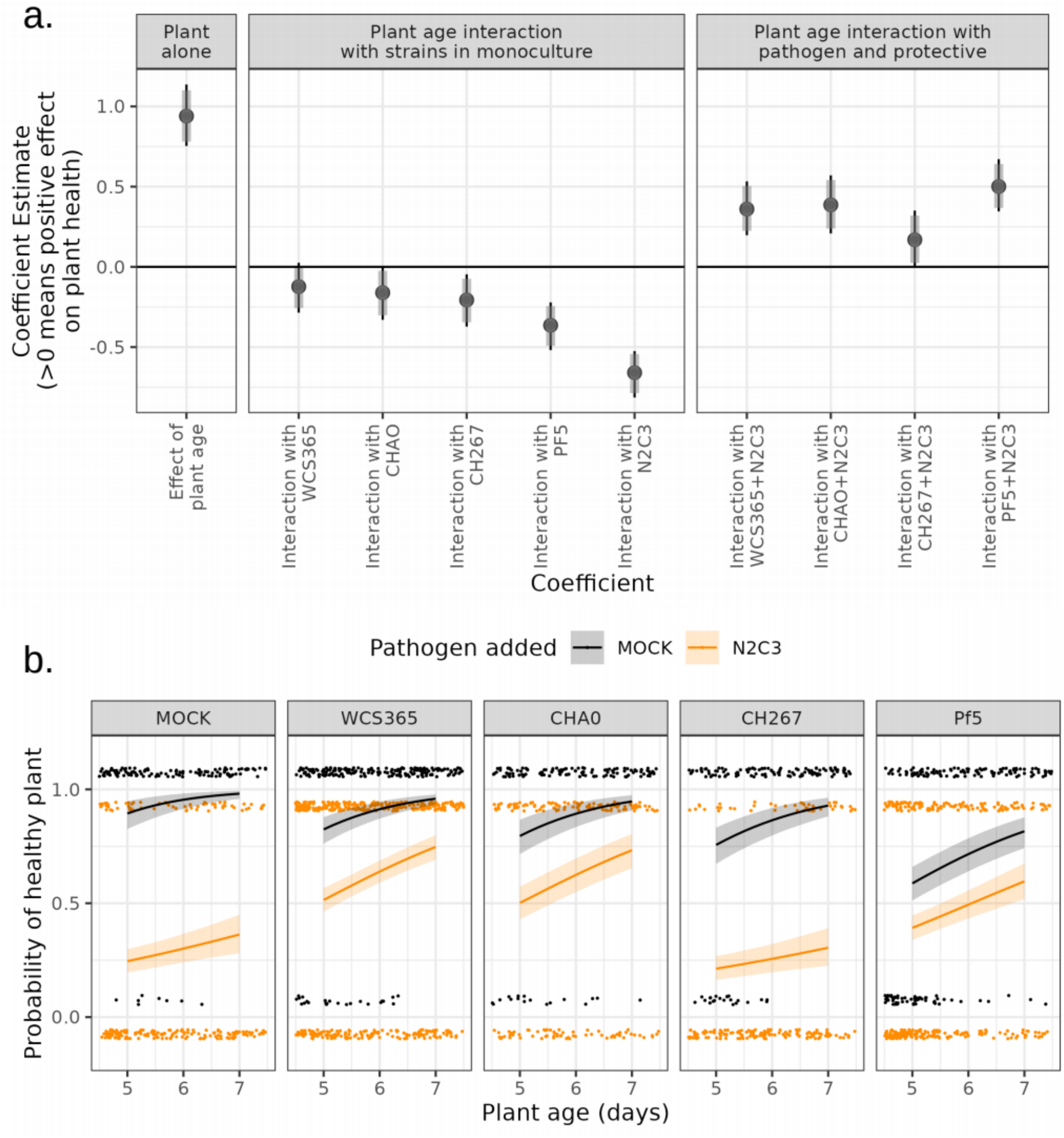


Figure S8. Older plants are less susceptible to infection and benefit more from protective symbionts than younger plants. (a) Coefficient estimates for different variables associated with plant age show that the effect of plant age interacts with bacterial strain. In the first panel, the coefficient for the independent (global) effect of plant age is positive, meaning older plants are more likely to remain healthy, overall. In the second panel, we show the interaction between age and each monoculture treatment. Negative coefficient numbers mean there is a weaker effect of age when treated with that strain, relative to the global positive effect of plant age. Here, a negative value for N2C3:plant age means that while plants are usually healthier with age, the effect of plant age when inoculated with N2C3 is less different between 5-day-old and 7-day-old plants (ie 7-day-old plants are not as healthy as one would predict if there was no interaction between N2C3 and plant age). In the third panel, we show the interaction of plant age and each N2C3 + commensal treatment. When coefficients are positive, it means older plants benefit more from protection of the commensal, relative to what might be expected due to plant age alone. (b) Using the model from (a), we generated prediction intervals for the probability of healthy plants. Dots represent single plant outcomes. Lines represent median posterior predictions according to the Bayesian Bernoulli model; ribbons represent 95% prediction intervals.

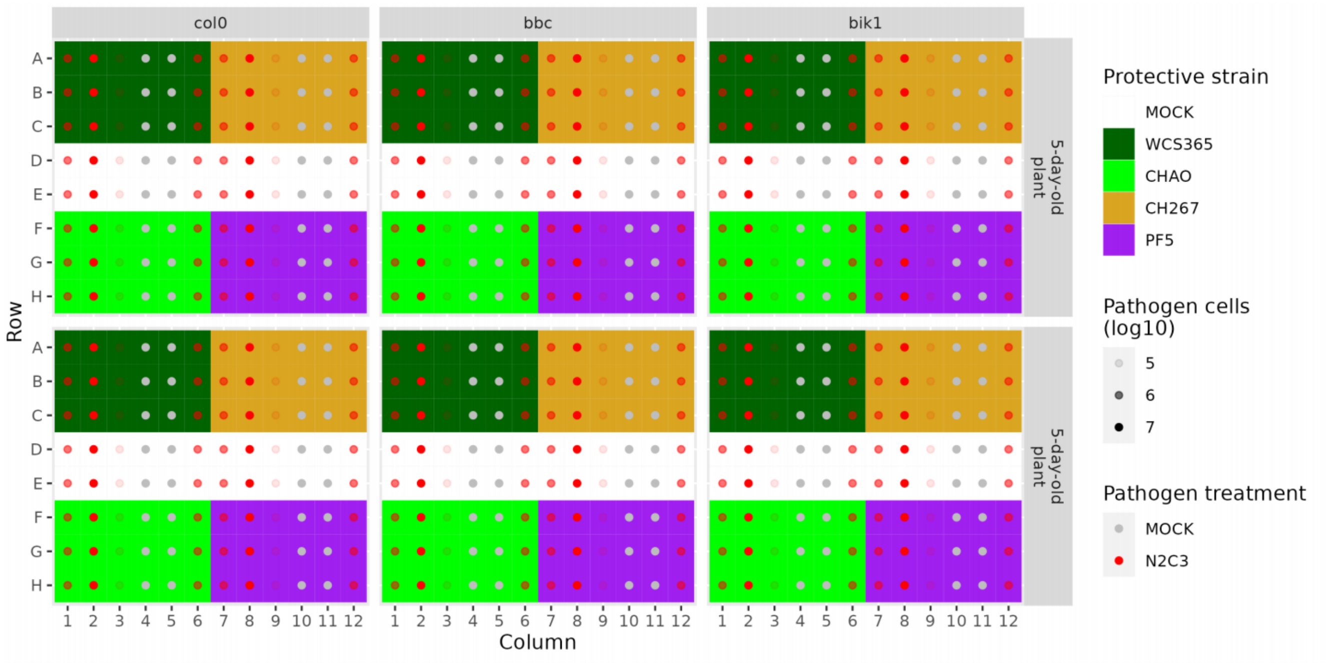


Figure S9. Plate map and experimental design for plant genotype experiment. Three plant genotypes (Col-0, *bbc*, and *bik1*) were inoculated with WCS365, CHA0, and Pf5 with or without N2C3 at three different protective:pathogen ratios. In our paper, we present only 1:1 and 1-0.1 protective:pathogen treatments because plants from 1:10 treatments were overwhelmingly “not healthy”, which means we could not observe any differences between control (Col-0) and treatment (*bbc*, *bik1*) plant genotypes.

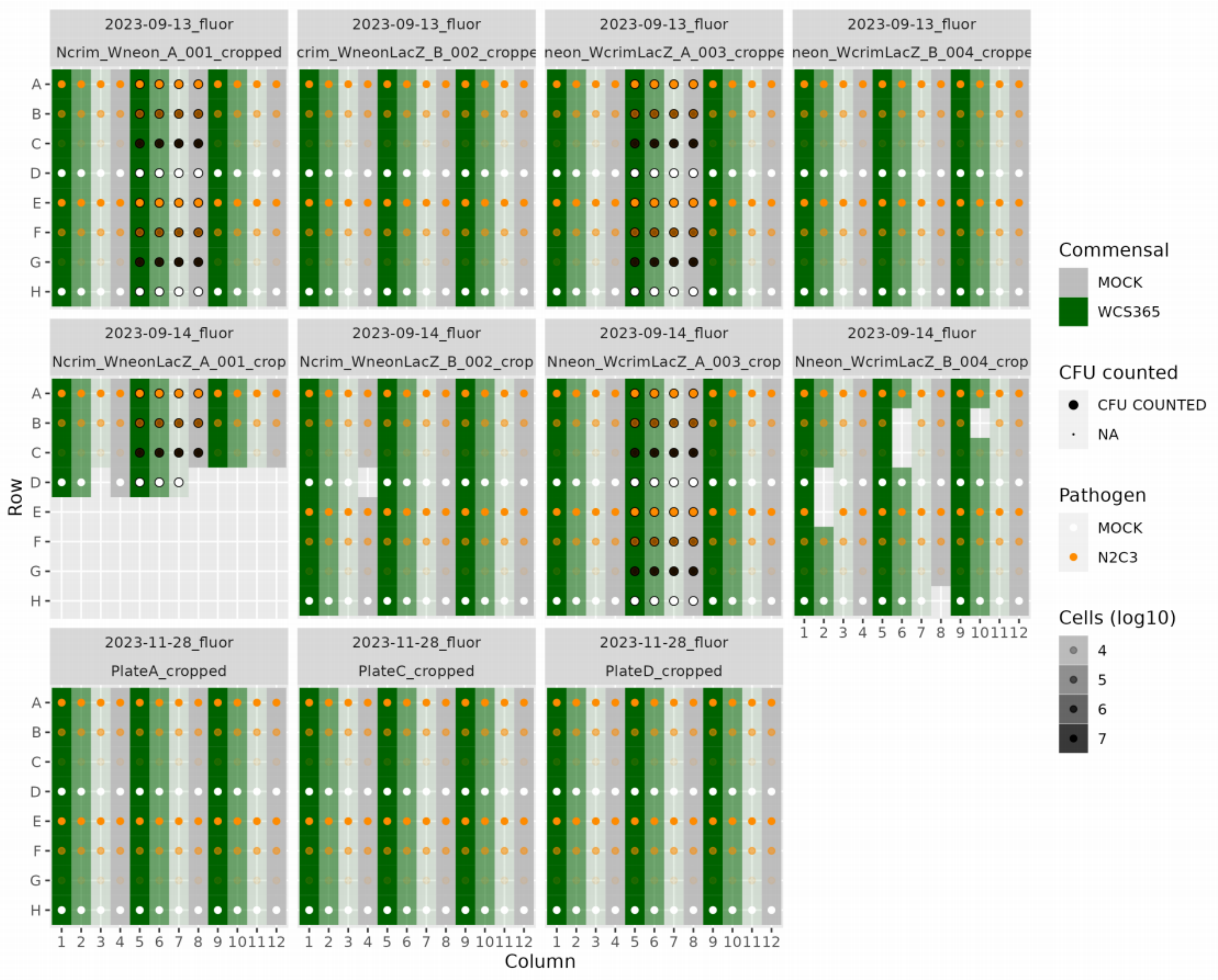


Figure S10. Plate map and experimental design for data shown in fluorescence experiment (Figure 4). Strains were labelled with either Crimson or Neon plasmids. Black outlined circles indicate wells that were used for serial dilution CFU counting, in order to get lacZ-based estimates of each strain’s cell density, which was fluorescence-independent.

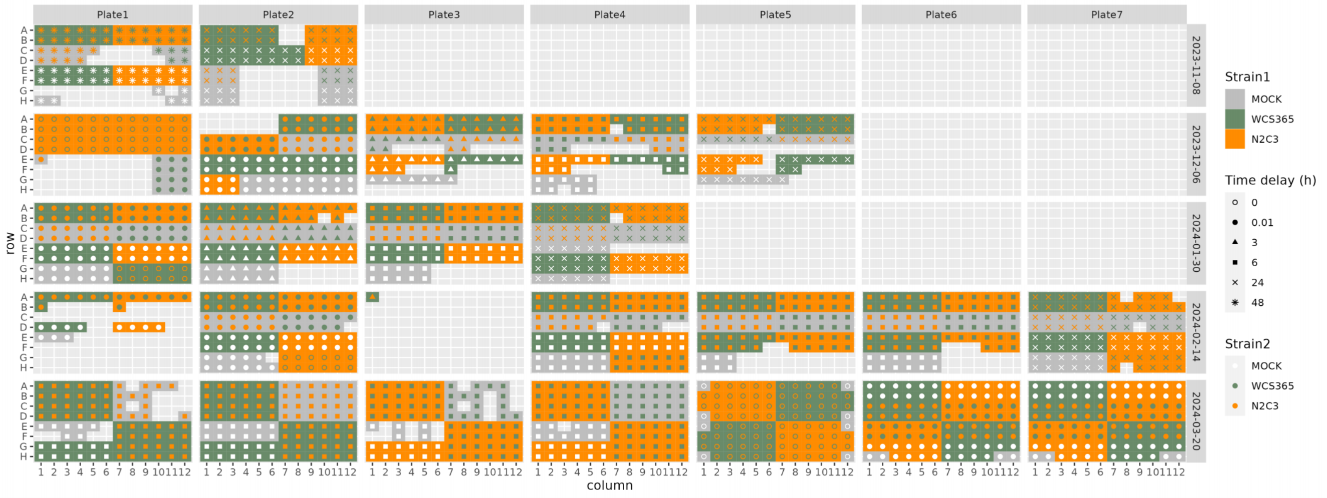


Figure S11. Plate map and experimental design for all priority effects experiment in Figures 5 and 6.

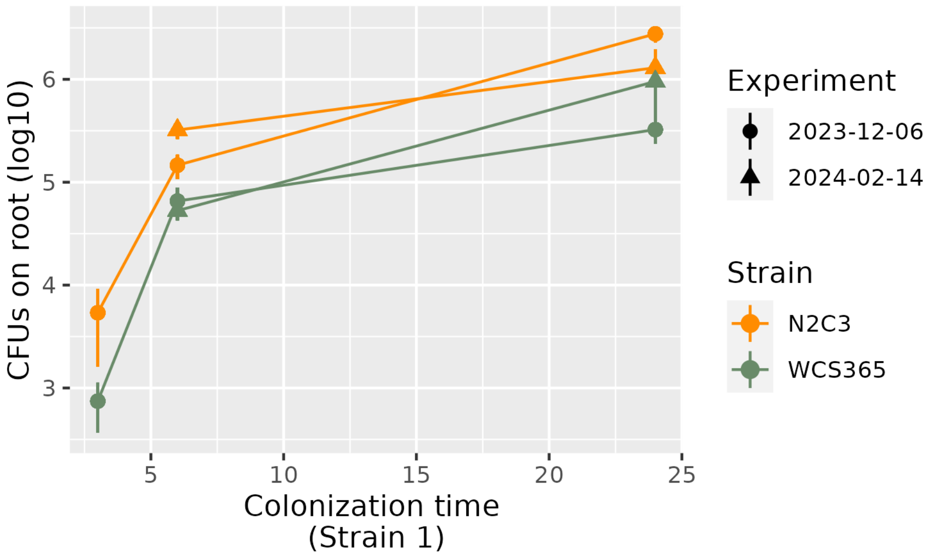


Figure S12. Cell densities of WCS365 and N2C3 on plant roots in monoculture across time. N2C3 reaches slightly higher cell densities than WCS365 on plant roots when in monoculture.

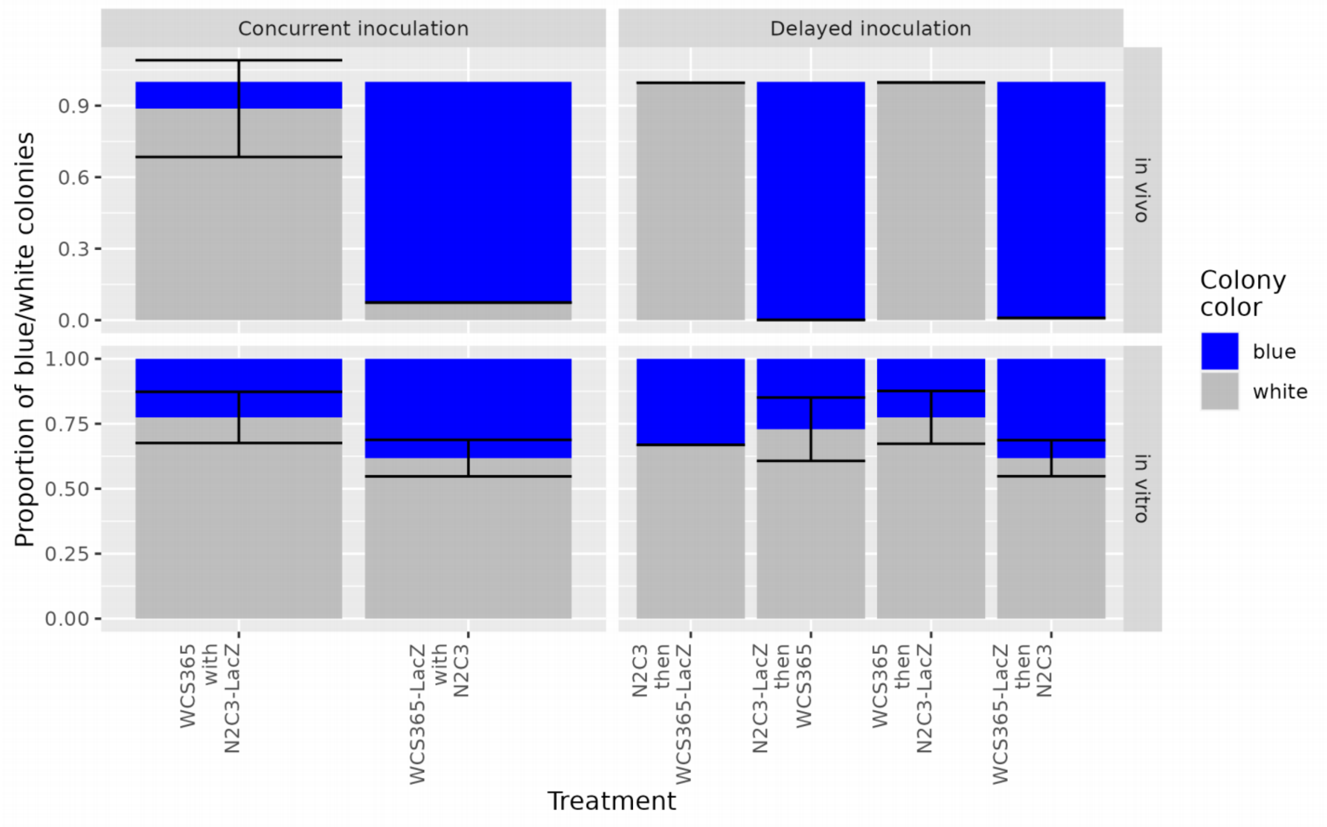


Figure S13. Strong priority effects exist between WCS365 and N2C3 when grown *in planta* but not *in vitro.* Shown are the proportions of WCS365 and N2C3 colonies on solid plant agar when grown with and without plants under different inoculation regimes. Plants were inoculated with WCS365 or N2C3 either at the same time (Concurrent inoculation) or with a 24 hour delay (Delayed inoculation).

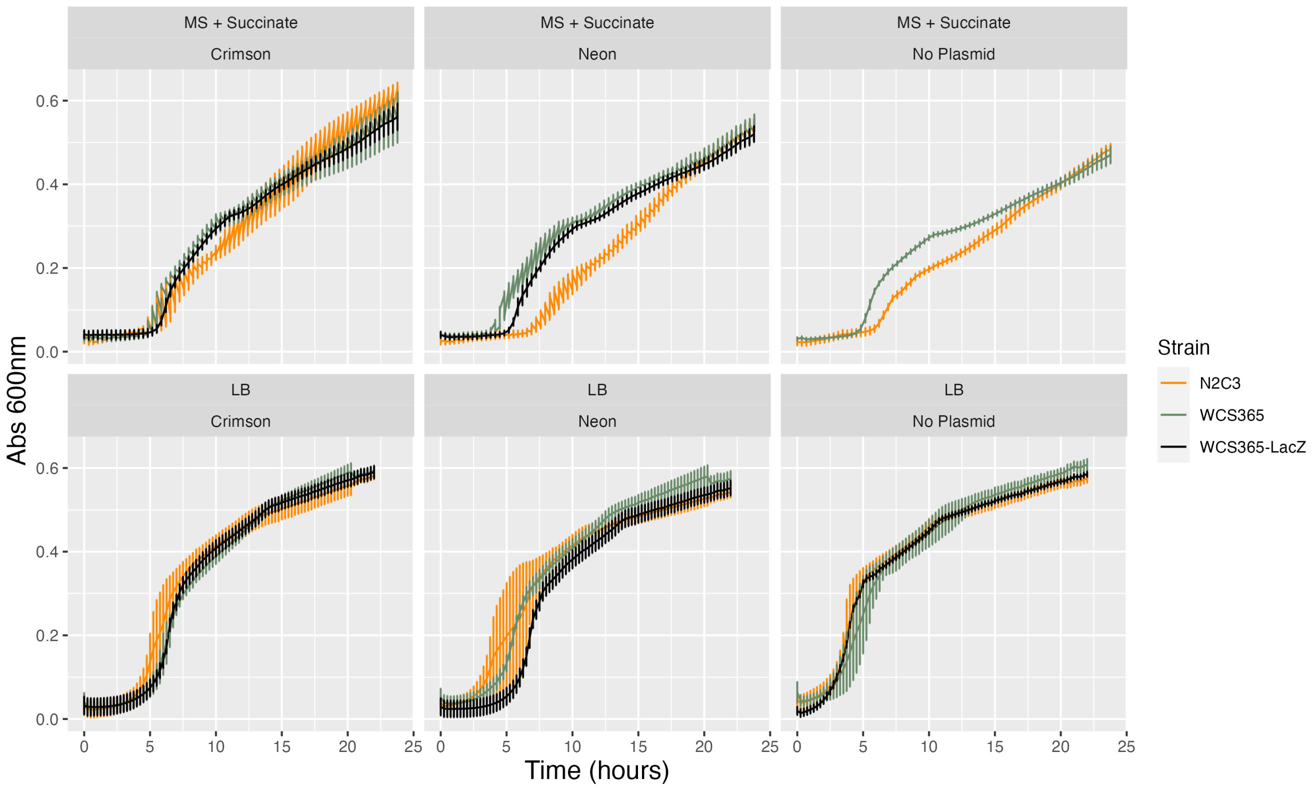


Figure S14. Growth curves of all strains used in experiments. WCS365 and N2C3 do not differ in growth rate or carrying capacity in LB media, with or without plasmids. There is a slight burden on N2C3 with both m-Crimson and m-Neon plasmids in minimal media.

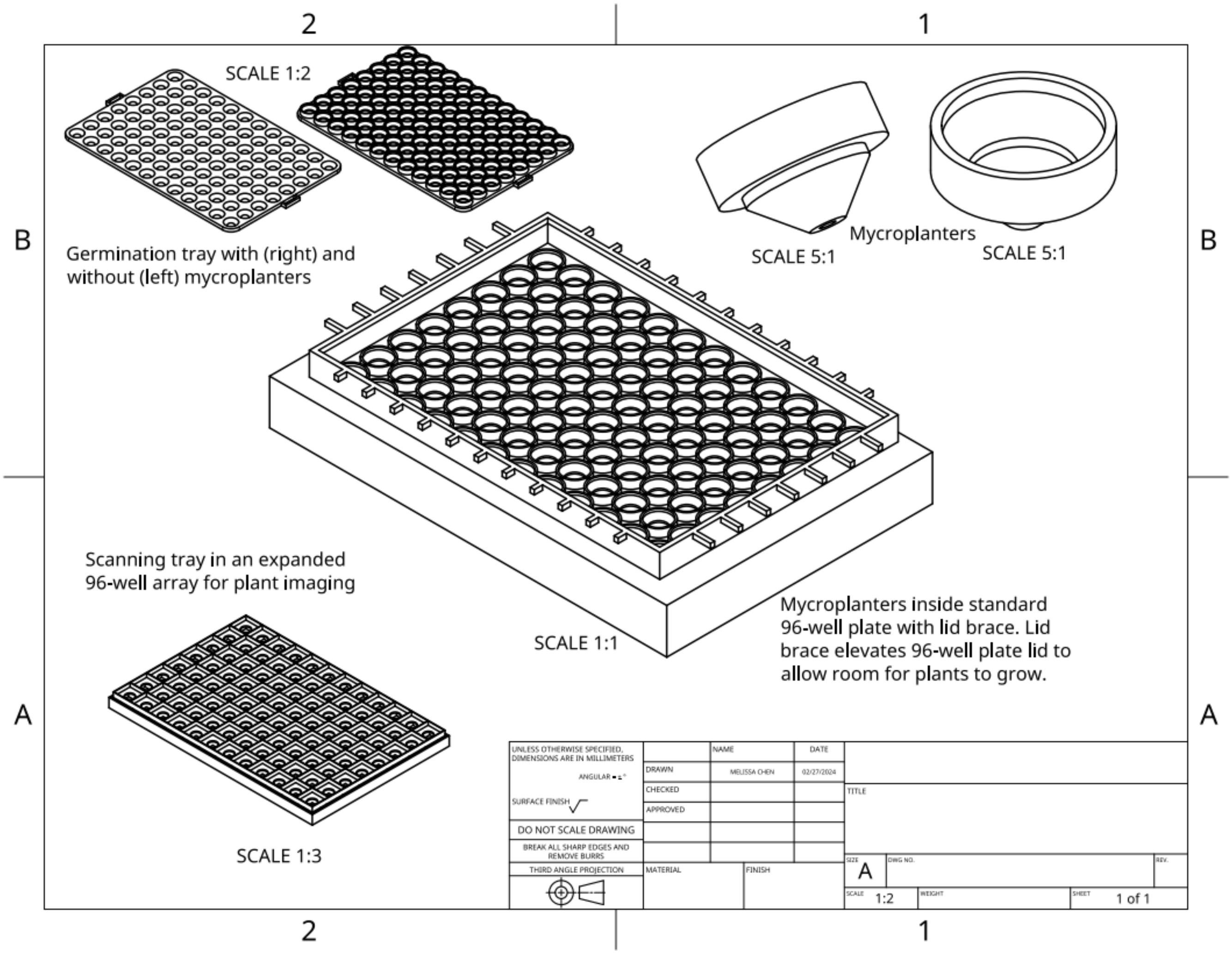


Figure S15. Drawings of all MYCroplanter components. Top left: germination tray used to hold MYCroplanter upright during seed germination on solid plant phytoagar. Top left: two individual MYCroplanter at different angles. Centre: MYCroplanters sitting in a standard 96-well plate with a lid brace. The lid brace elevates the 96-well plate lid (which comes with the standard sterile 96-well plate) so that plants have room to grow upward. Bottom left: scanning tray in a 96-well array was used to image plants because plants in the 96-well array grow too close together. The scanning tray is made up of an upper tray (which holds the mycroplanter) and a bottom tray (to prevent roots from touching common surfaces), and its array is wider than that of a standard 96-well plate, which means leaves are spread out and do not touch during imaging. To image plants, a 20.32 x 27.94 cm (8 x 11 inches) piece of glass (not shown) is placed ontop of the plants and the entire unit is flipped upside down and scanned using a printer scanner.

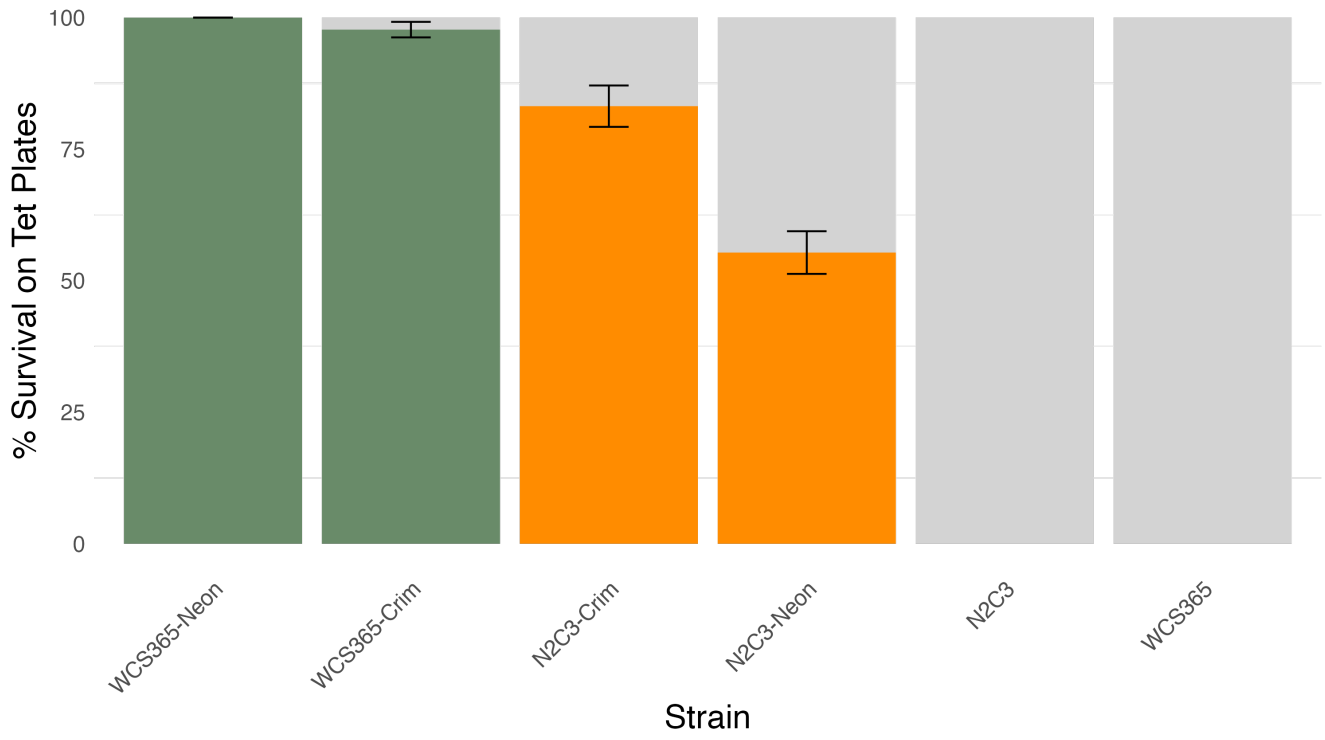


Figure S16. Plasmid loss for WCS365 and N2C3 strains with Neon and Crimson plasmids. N2C3 strains lost both plasmids at higher rates than WCS365, and this effect was particularly strong for N2C3-Neon.
