## Supplemental methods for "Order among chaos: high throughput MYCroplanters can distinguish interacting drivers of host infection in a highly stochastic system"

**SUPPLEMENTARY METHODS**

The $\log_2$ fold change of blanked fluorescence values was used to estimate relative abundance of fluorescent strains. The mathematical logic for our method is as follows:

The cell density of a culture (OD) can be estimated from fluorescence values (F) using a standard curve where the relationship is described by:

$$OD = b + m*F$$

F is the raw fluorescence value and OD is the corresponding OD600 of a bacterial culture. In a perfect standard curve, the intercept term (b) approaches zero because no fluorescence should indicate no cells (after blanking). Therefore, the equation relating OD to fluorescence can be simplified to:

$$OD = m*F$$

The relative abundances of two strains in a mixed culture can be described using log2 fold-change. Thus, we can calculate the log2 fold-change of two strains using the following formula:

$$\log_2 ( OD_{crim} / OD_{neon} ) = \log2( m_{crim}*F_{crim} / m_{neon} *F_{neon})$$

where $m_{crim}$ and $m_{neon}$ are the slopes from a standard curve relating OD to fluorescence, and $F_{crim}$ and $F_{neon}$ are the raw fluorescence values measured from each strain. The ratio of ODcrim to ODneon is therefore the log2 fold-change ratios of each strain, based on their fluorescent ratios. Through log identities, this equation can be simplified to:

$$\log_2 ( OD_{crim} / OD_{neon} ) = \log2( m_{crim}/ m_{neon} ) + \log2( F_{crim} /F_{neon})$$

The $\log_2$ fold change of fluorescence is equal to the $\log_2$ fold change of cell densities, plus some constant (the $\log_2$ ratio of slopes). In our approach, we directly estimate the $\log_2$ ratio of slopes constant by conducting lacZ cell counts of WCS365 and N2C3 in a subset of treatments, and using these real cell counts to adjust the y-intercept for the $\log_2$ fold change ratio line fit.

Thus, the $\log_2$ fold change between competing strains was calculated by first calculating $\log_2$ fold change of blanked fluorescence values, and then linearly transforming those values by a constant which we derived by comparing lacZ plate counts
